# Diploid potato lines for the study and improvement of starch metabolism and structure

**DOI:** 10.64898/2025.12.19.695480

**Authors:** Thomas Navarro, Yordan Dolaptchiev, Oluwaseyi Bello, Ciara O’Brien, Andrés Ortiz, Jonathan D. G. Jones, David Seung

**Affiliations:** John Innes Centre, Norwich Research Park, Norwich, NR4 7UH, UK; The Sainsbury Laboratory, University of East Anglia, Norwich, NR4 7UH, UK

## Abstract

Diploid potato breeding enables faster genetic improvement via selection against deleterious alleles in inbred lines, unlike breeding by intercrossing tetraploid varieties. Starch is the major source of calories in potato tubers, but the starch properties of diploid lines have rarely been reported. In this study, we provide a comprehensive characterisation of tuber and starch properties in two diploid lines that are early isolates from the Solynta breeding program, B26 and B100, and their F_1_ hybrids. B100 produced fewer, but larger tubers compared to B26, and both diploid lines produced tubers that are smaller than the tetraploid variety, Clearwater Russet. The low tuber yield of B100 correlates with its high self-compatibility and fruit production. Pruning of fruits in B100 significantly increased total tuber yield per plant by stimulating more tuber initiations, but had no effect on average tuber weight, starch content or starch structure. Among the diploid, hybrid and tetraploid lines examined, there were no differences in the total starch content of tubers. Although amylopectin structure and amylose content were similar between the two diploid lines and the tetraploid comparison, B26 had elevated levels of resistant starch and a striking elongated granule morphology. Our results showcase the variation in source-sink relations and starch structure in diploid potato breeding material, demonstrating their potential for research into the genetics underpinning metabolic and quality traits.

## INTRODUCTION

Potato (*Solanum tuberosum L.*) is the fourth most produced food crop globally, and the most produced non-cereal crop (FAOSTAT, 2023). The major source of calories in potato is starch, the main carbohydrate component of the tubers. Starch is important for nutritional value as well as functional applications due to its unique physicochemical properties (Ashraf et al., 2019; Li et al., 2019; Zeeman et al., 2010). It is mainly composed by two glucose polymers, amylopectin and amylose, both made of α-1,4 linear chains and α-1,6-linked branched points. Amylopectin is the major polymer constituent, typically 67-80% of potato starch, and is highly branched. Amylose comprises the remaining 20-33%, and is primarily linear (Dupuis & Liu, 2019; Seung, 2020). Starch forms semi-crystalline, insoluble granules, which for potato have an average diameter of 40-50 µm, but ranging from 10-110 µm (Waterschoot et al., 2015). Both polymer structure and granule morphology influence the functional and nutritional properties of starch (Chen et al., 2021).

Most commercial potato cultivars are tetraploid, highly heterozygous, and self-incompatible. Their vegetative propagation through genetically identical seed tubers results in long generation cycles and pathogen contamination (Lindhout et al., 2018; Vos et al., 2015). Tetraploid potato breeding involves intercrossing varieties and screening for rare F_1_ progeny that are better than either parent, and results in low genetic gain. To tackle these challenges, significant progress has been made in developing homozygous inbred diploid lines for F_1_ hybrid seed production, accelerating genetic gain and enabling selection against deleterious alleles (Bethke et al., 2022; Lindhout et al., 2011; Zhang et al., 2021). Hybrid breeding has proven successful in crops like maize and could revolutionise potato farming by enabling the use of true potato seeds, offering genetic stability, reduced storage costs, and lower disease risks compared to seed tubers (Stokstad, 2019). Additionally, the simplified genetic background and crossability of diploid lines provide an efficient platform for studying the genetics of important traits and enable elimination of transgenes required for gene editing.

Despite their potential in potato breeding and biotechnology, there are no reports of starch traits in this novel diploid germplasm. It is therefore not known whether there is variation in key starch parameters within modern diploid breeding programs in comparison to extensively characterised tetraploid varieties. Here, we characterised the starch properties of two inbred lines from the early stages of Solynta’s diploid breeding program, B26 and B100, and two hybrids generated between these lines. Although a complete pedigree is not available, these diploid lines were derived from crosses between the diploid wild potato *Solanum chacoense* and dihaploid lines produced from tetraploid varieties (Hutten et al., 1995; Lindhout et al., 2018). We compared these lines to the commercial tetraploid, Clearwater Russet. While Clearwater Russet had characteristics that were typical for previously characterised tetraploid potato lines, the two diploid lines had several distinct starch and physiological features. This variation can be exploited for the further study of metabolic traits, and the lines represent a promising platform for potato biotechnology.

## RESULTS

### Plant biomass and tuber yield varied among diploid lines

To assess the phenotype of the two diploid lines (B26 and B100) and their F_1_ hybrids (YD2 and YD3), plants were grown under glasshouse conditions. The two parent lines differed in growth habit, with B100 plants generating many more flowers and berries than B26 (Figure 1A). This was at least partially due to flower bud abortion in B26 (Figure S1). However, B26 had visibly more tubers than B100 (Figure 1B). Tubers were harvested from 17-week-old plants (Figure 1C), and shoot biomass was quantified as the total fresh weight of above-ground tissues immediately prior to tuber harvest. B100 had significantly higher shoot biomass than B26, and comparable biomass to the commercial tetraploid, Clearwater Russet (Figure 1D). The shoot biomass of the two hybrids, YD2 and YD3, was significantly higher than either diploid parent.

**Figure 1:**
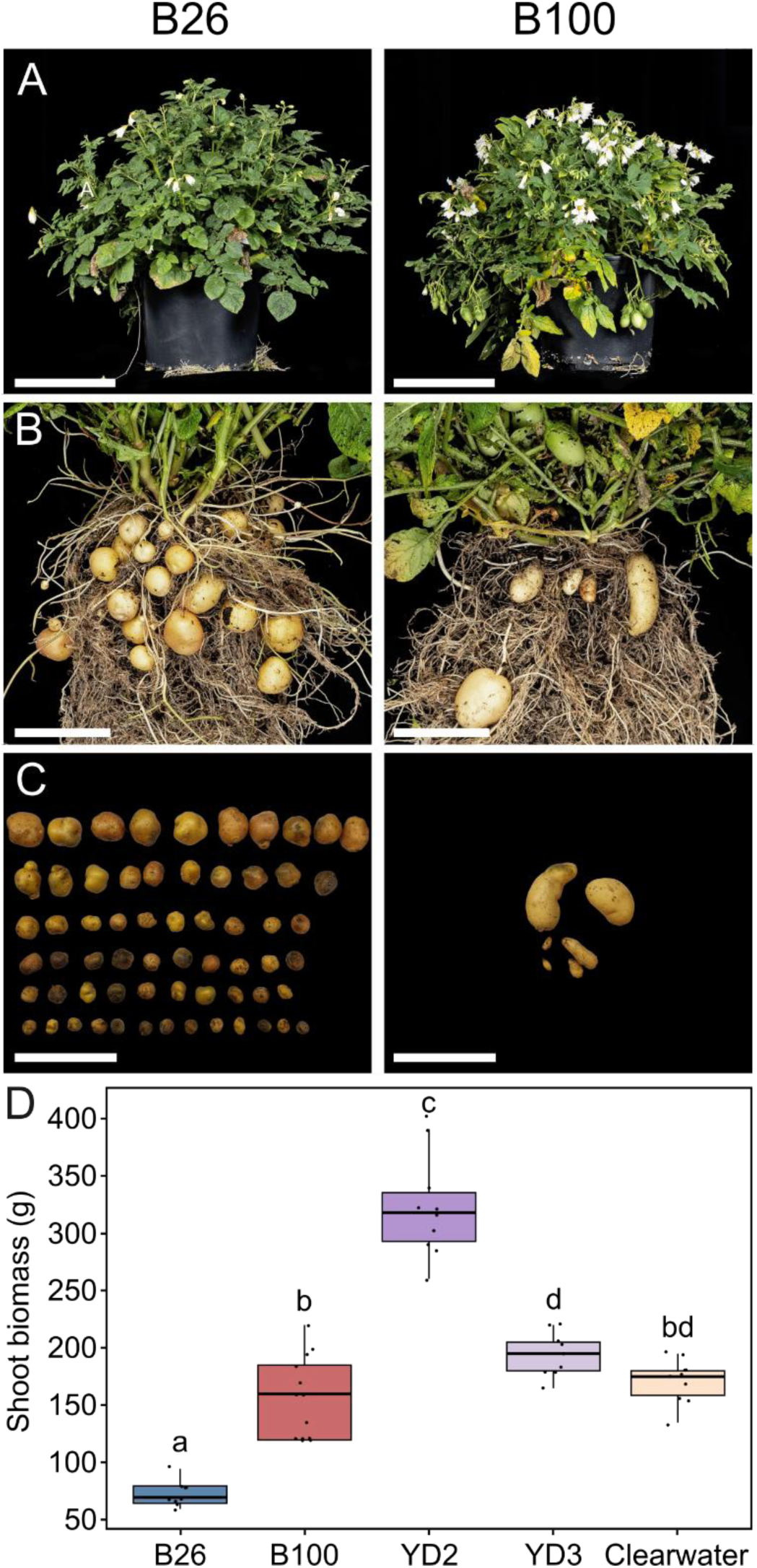
Growth and tuber phenotypes of diploid potato lines. **(A)** Photographs of a representative 12-week-old plants of lines B26 and B100. Bar = 15 cm. **(B)** Same plant as panel A, showing tubers after washing away the soil. Bar = 5 cm. **(C)** Tubers from a representative plant after harvesting (at 17 weeks). Bar = 10 cm. **(D)** Shoot biomass as fresh weight of total above-ground biomass measured before harvest (at 17 weeks) from diploid lines (B26, B100), diploid hybrids (YD2, YD3), and commercial tetraploid variety (Clearwater Russet). For the boxplots, the band inside the box represents the median, while the top and the bottom of the box represent the lower and upper quartiles, respectively. The whiskers represent values within 1.5× of the interquartile range. Dots represent individual data points (n = 9-13 per genotype), with each point representing an individual plant. Values with different letters are significantly different from each other under a one-way ANOVA with Tukey’s posthoc test (p <0.05).

After harvest, we quantified tuber traits, which again varied between B26 and B100. Overall, B100 produced significantly lower total tuber yield per plant than B26, quantified as the total weight of all tubers harvested per plant (Figure 2A). Tuber yield per plant in B26 was comparable to Clearwater Russet. However, B26 produced extremely large numbers of tubers, with a mean of 72 tubers per plant, whereas B100 produced at most 10 tubers per plant (Figure 2B). The mean number of tubers per plant was not significantly different between B100 and Clearwater Russet. The total tuber yield and tuber number for YD2 and YD3 were intermediate between the parent lines. Although there were no significant differences in the average weight of the diploid tubers, which was likely due to the high variability in this trait between plants. The average weight of the tubers in B26 was consistently low, between 3.2–5.0 g among the 9 replicate plants (Figure 2C). This is in strong contrast to B100, where average tuber weight was more variable between replicates, from 0.7 g and up to 20.3 g. Clearwater Russet consistently produced large tubers, with an average tuber weight that was 20.5–86.8 g. B26 and B100 are therefore distinct in tuber forming habit, where B26 consistently produces many small tubers, while B100 tubers are infrequent but can be larger than those of B26. However, none of the diploid lines produced tubers that were comparable in size to Clearwater Russet.

**Figure 2:**
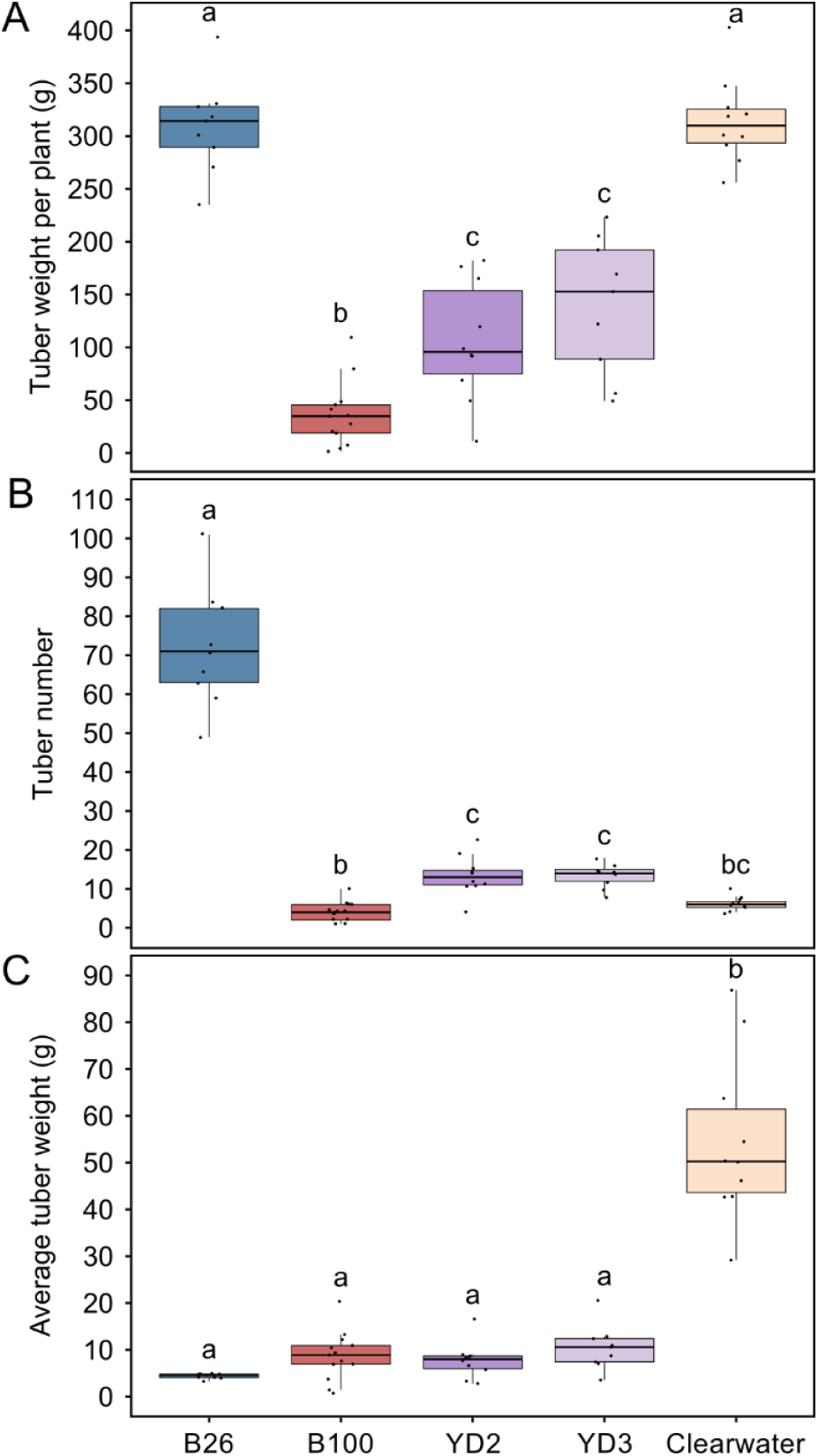
Tuber yield measurements of diploid potato lines. Tubers were harvested from 17-week-old plants of diploid lines (B26, B100), diploid hybrids (YD2, YD3), and commercial tetraploid (Clearwater Russet). **(A)** Total tuber weight harvested per plant. **(B)** Number of tubers per plant. **(C)** Average weight of individual tubers, calculated for each plant. For the boxplots, the band inside the box represents the median, while the top and the bottom of the box represent the lower and upper quartiles, respectively. The whiskers represent values within 1.5× of the interquartile range. Dots represent individual data points (n = 9-13 per genotype, with each point representing an individual plant). Values with different letters are significantly different from each other under a one-way ANOVA with Tukey’s posthoc test (p <0.05).

### Berry formation in B100 competes with tubers as a carbon sink

In addition to the low tuber yield observed in B100, a striking feature of B100 plants was extensive flowering and fruit set, reflecting a high degree of self-compatibility in the line (Figure 3A). The berries resembled an elongated tomato fruit (∼3 cm long), and iodine staining in the immature berry revealed starch accumulation in the locular cavity surrounding the seeds (Figure 3B). We therefore hypothesised that the berries could act as a competing carbon sink during tuber development, contributing to the low tuber yield of B100. To address this, we mechanically disrupted the flowers in a subset of B100 plants to prevent berry formation and compared shoot and tuber responses to control plants where flowering and berry formation was left intact.

**Figure 3:**
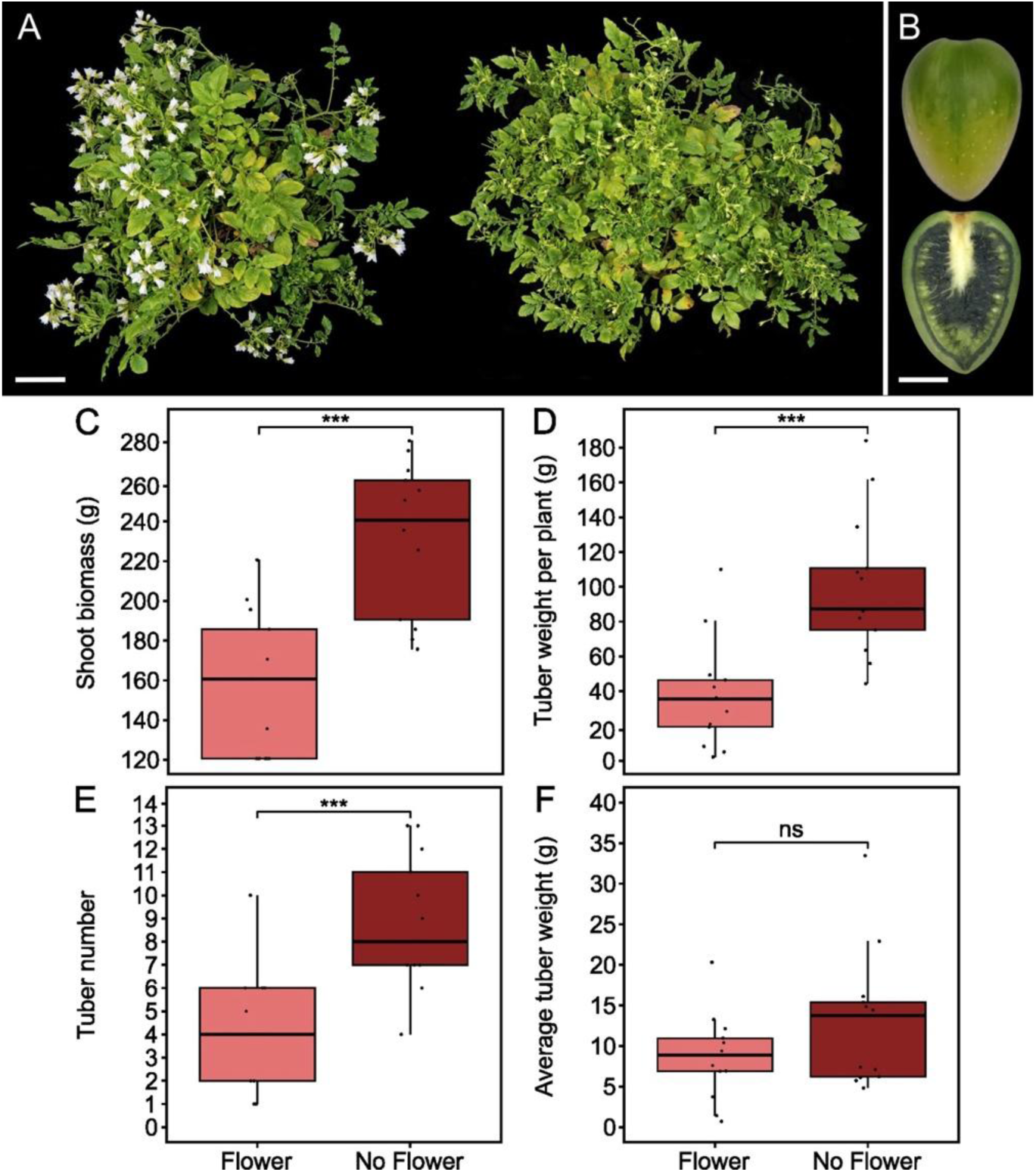
Flower removal in B100 increases shoot biomass and number of tubers. **(A)** Photographs of two representative 12-week-old plants from B100 without (left) and with (right) flower removal. Bar = 10 cm. **(B)** Photograph of a harvested berry from B100. Top half shows a whole berry, while the bottom shows a longitudinal section stained with iodine to visualise starch (stained dark). Bar = 1 cm. **(C)** Shoot biomass as fresh weight of total above-ground biomass measured before harvest (at 17 weeks) in plants with and without flower removal. **(D)** Total tuber weight harvested per plant. **(E)** Number of tubers per plant. **(F)** Average weight of individual tubers, calculated for each plant. For the boxplots, the band inside the box represents the median, while the top and the bottom of the box represent the lower and upper quartiles, respectively. The whiskers represent values within 1.5× of the interquartile range. Dots represent individual data points (n = 13 per genotype, with each point representing an individual plant). *** represents significance under a two-tailed t-test at p < 0.001. ns = not significant.

Flower disruption led to altered plant development, increasing production of stems and leaves. This led to 43% higher shoot biomass in plants with flower disruption compared to non-disrupted plants at the time of harvest (Figure 3C). Total tuber weight per plant was significantly higher in plants where berry formation was prevented in comparison to control plants with berries (Figure 3D). These findings confirm that berries represent a competing sink, and disrupting their formation allows greater partitioning of carbon towards tubers and shoot biomass. Interestingly, the increased tuber yield per plant after flower disruption was primarily driven by an increase in tuber initiation. Plants with berries removed had almost double the number of tubers compared to control plants (Figure 3E). There was no significant effect on the average tuber weight after flower disruption (Figure 3F).

Taken together, B26 and B100 differ in growth habit in that B26 yields many tubers, while B100 yields very few. The large amount of fruit formation in B100 is one of the reasons for its low tuber yield. The diploid lines therefore offer physiological variation that is valuable for studying source-sink relations, in addition to berry and tuber initiation.

### Starch composition and structure vary among the diploid lines

Since starch properties have not been characterised in diploid potato lines, we aimed to quantify major starch parameters between B26 and B100 and its hybrids. We first quantified dry matter content and total starch content in the lines. Although there were some significant differences between the lines in dry matter content, the differences were relatively small (< 5%), and there were no significant differences in total starch content. However, when we quantified the amount of resistant starch, which is an important starch parameter for tuber nutritional properties, B26 had significantly more resistant starch than the other lines (15.0% increase relative to B100, 24.7% increase relative to Clearwater Russet). As increases in resistant starch can occur due to increases in the amount of amylose, we quantified amylose content (Figure 4D). The average amylose content ranged from 23.2% to 25.4% in the diploids and hybrids, compared to 21.5% in Clearwater. However, there were no significant differences between the lines. Therefore, the increased levels of resistant starch in B26 are likely due to starch properties other than amylose content.

**Figure 4:**
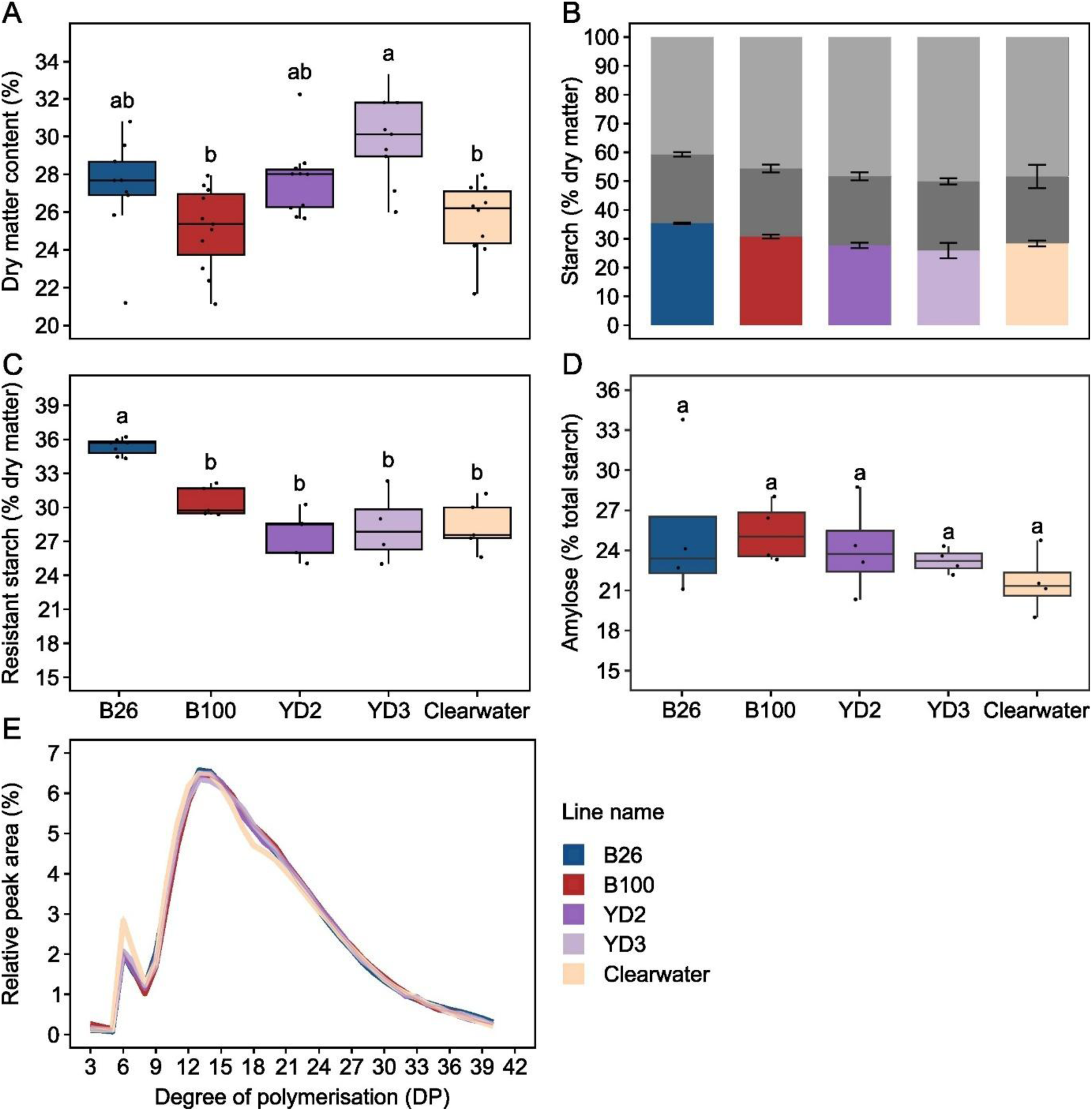
B26 has higher levels of resistant starch compared to the other diploid lines. **(A)** Percentage of dry matter in potato tubers in diploid lines (B26, B100), diploid hybrids (YD2, YD3), and commercial tetraploid (Clearwater Russet). Dots represent individual data points (n = 9-11 per genotype, with each point representing an individual plant). **(B)** Total starch content extracted from the dry matter fraction of potato tubers. Stacking of bars represents the different fractions of starch with the bottom chart in colours representing resistant starch, the middle dark grey chart for non-resistant starch, and the top light grey chart for non-starch fraction. **(C)** Boxplot of resistant starch fraction in potato dry matter from panel B. Dots represent individual data points (n = 4-5 per genotype, with each point representing tubers harvested from an individual plant). **(D)** Amylose content as a percentage of total starch from potato tuber (n = 4 per genotype, with each point representing a different plant). **(E)** Amylopectin chain length distribution determined using High Performance Anion Exchange Chromatography with Pulsed Amperometric Detection (HPAEC-PAD). The relative peak values for each DP, with a solid line corresponding to the mean value, and the shaded area for the standard error. For all boxplots in this figure, the band inside the box represents the median, while the top and the bottom of the box represent the lower and upper quartiles, respectively. The whiskers represent values within 1.5× of the interquartile range. Values with different letters are significantly different from each other under a one-way ANOVA with Tukey’s posthoc test (p <0.05).

To assess for other changes in polymer composition and structure, we determined the amylopectin chain length distribution using High Performance Anion Exchange Chromatography with Pulsed Amperometric Detection (HPAEC-PAD). There were no differences in amylopectin chain length structure between the diploid lines. However, all the diploid lines had an altered chain length distribution compared to the tetraploid, with a lower proportion of short chains (DP 6 & 7) and increased proportion of medium chains (DP 16-21) (Figure S2B and S2C).

Finally, we examined the morphology of the starch granules. We first quantified starch granule size distributions by analysing purified starch granules on the Coulter counter. Potato starch granules are typically elliptic in shape (Hochmuth et al., 2025; Jane et al., 1994). Using Scanning Electron Microscopy (SEM) on purified starch granules, we observed that B100 and Clearwater Russet showed this typical granule morphology of potato starch (Figure 5A). Surprisingly, B26 starch granules showed a distinct morphology, where the granules were more elongated. To our knowledge, such a granule morphology has never been reported for storage starches.

**Figure 5:**
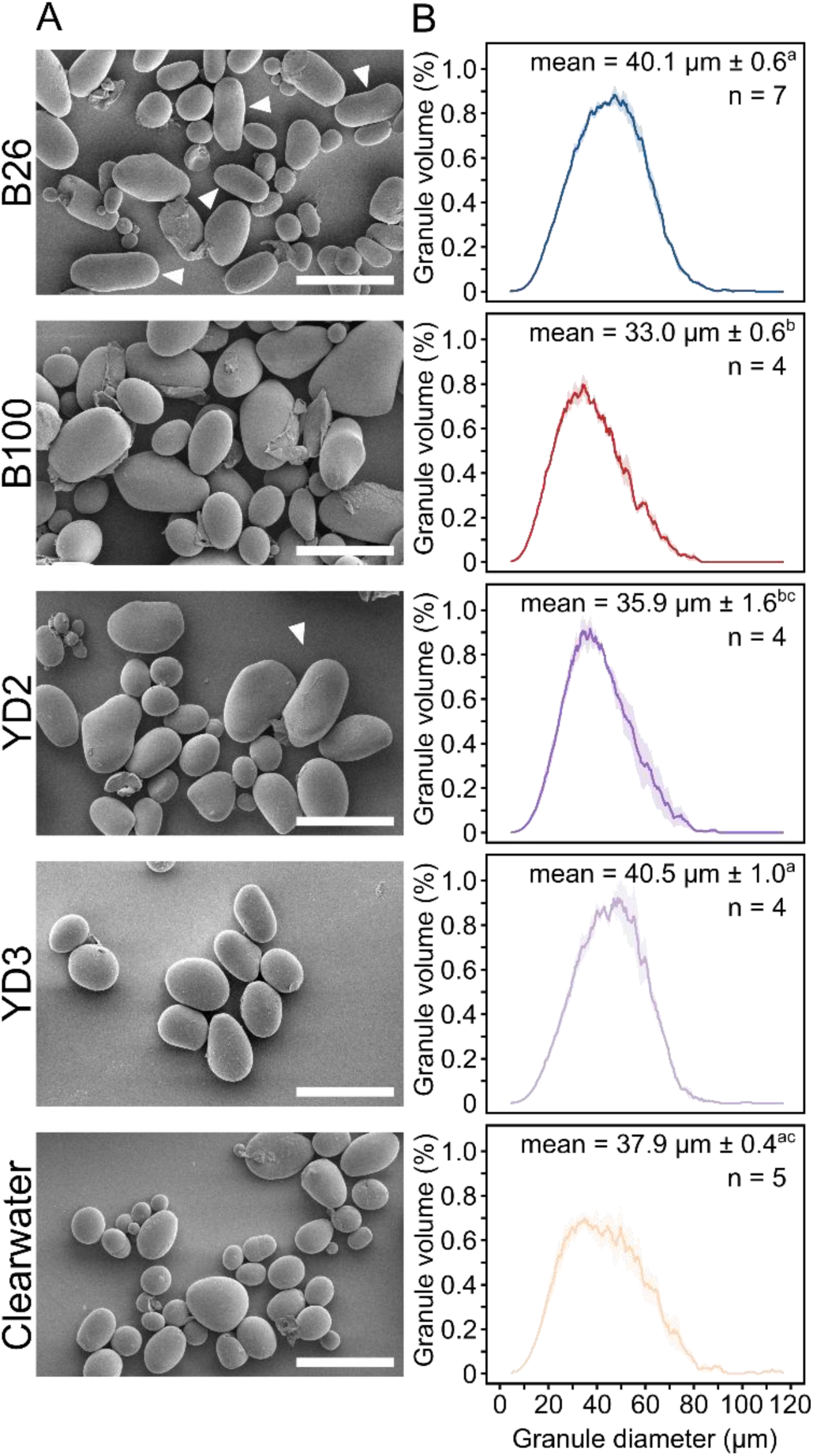
B26 starch granules are large and elongated. **(A)** Scanning Electron Microscopy (SEM) of purified starch granules from diploid lines (B26, B100), diploid hybrids (YD2, YD3), and commercial tetraploid (Clearwater Russet). Arrows highlight elongated starch granules in B26 and YD2. Bar = 50 μm. **(B)** Starch granule size distribution plots showing percentage of relative granule volume against granule diameter. The line represents the mean distribution from n = 4-7 plants per genotype, while shading represents the standard error. The mean granule size ± standard error for each genotype is indicated on each panel. Values with different letters are significantly different from each other under a one-way ANOVA with Tukey’s posthoc test (p <0.05).

For a quantitative assessment of starch granule morphology, we used particle size analysers. Firstly, we used the Coulter counter to quantify granule size distributions by measuring particle volume (Figure 5B). The granules followed a unimodal distribution in all genotypes. However, B26 had significantly larger mean granule size than both B100 and Clearwater Russet. For the hybrids, the mean granule size of YD2 resembled that of B100, while YD3 resembled that of B26. Secondly, we used the QICPIC analyser, which uses imaging to quantify granule shape parameters. The degree of elongation in starch granules can be quantified using the aspect ratio, which is the length of the minor axis divided by that of the major axis, such that the aspect ratio decreases from 1 as granules become more elongated (Hochmuth et al., 2025). All lines showed a broad distribution of aspect ratios. However, as we expected, the distribution of aspect ratio was similar between B100 and Clearwater Russet, but shifted towards lower aspect ratios in B26, suggesting that the granules in B26 are more elongated. Interestingly, YD2 followed the same pattern as B26, while YD3 behaved like B100 (Figure 6). The most likely explanation is that the elongated granule trait in B26 is regulated by a single dominant, heterozygous allele that segregated in the hybrid progeny.

**Figure 6:**
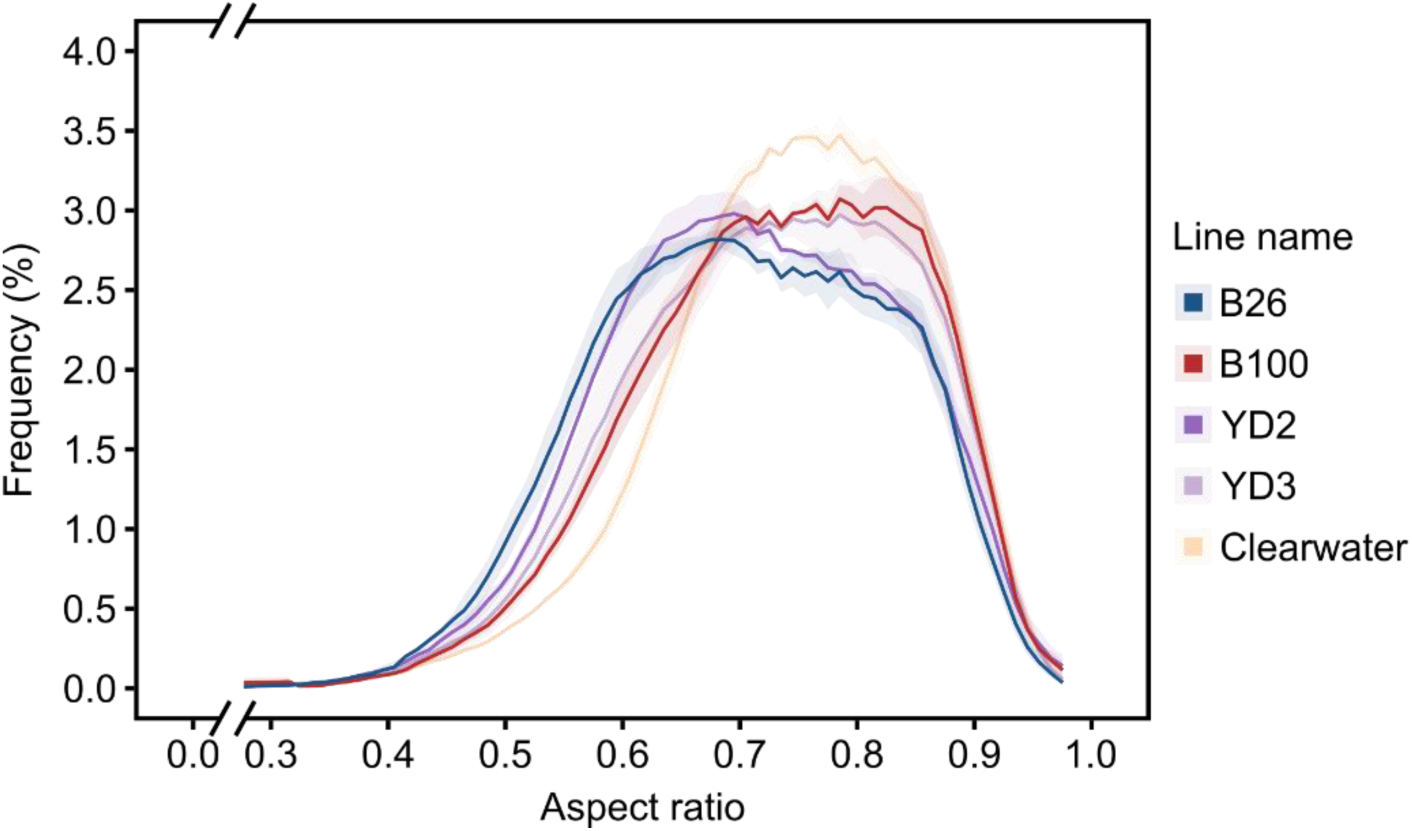
Elongated starch granule phenotype segregated in YD2 and YD3 hybrids. QICPIC particle shape analysis of diploid lines B26, B100, YD2 and YD3, and Clearwater Russet as a tetraploid comparison. The frequency of events is plotted against the aspect ratio (0 < ψ_A_≤ 1) with the solid line corresponding to the mean value, and the shaded area for the standard error (n = 3 replicate starch extractions, each from tubers harvested from separate plants).

## DISCUSSION

In this study, we demonstrated that two diploid potato lines, B26 and B100, have diverse physiological and tuber starch characteristics. The two lines contrasted each other in traits such as tuber number, fruit set, resistant starch content, and starch granule morphology (Figures 2, 4, 5, and 6). Although these lines still have some level of heterozygosity, compared to commercial tetraploids they have less allelic variation, a smaller genome size, and can be crossed (Lindhout et al., 2018). This genetic amenability, combined with the observed phenotypic variation, make them great models for genetic research on physiological and tuber quality traits, and a source of valuable alleles for breeding.

### Diverse tuber traits in diploid potato as a resource to study tuberisation

B26 and B100 had distinct growth habit and tuber phenotypes, despite coming from the same breeding program. Tuber size in all diploid lines was much smaller than the commercial tetraploid, Clearwater Russet (Figure 2C). However, the number of tubers in B26 was strikingly high compared to the other lines, and although each tuber was relatively small, the total tuber yield per plant (by weight) was comparable to Clearwater Russet (Figure 2A). This suggests that tuber initiation is more active in B26 compared to the other lines. By contrast, B100 had the lowest number of tubers and total tuber yield per plant among the lines. We demonstrated that this was partially due to the high level of flowering and fruit set in this line, since removing flowers doubled the total tuber yield per plant (Figure 3D).

Interestingly, the increase in tuber yield in B100 after flower removal was primarily due to an effect on tuber initiation. Flower removal doubled the number of tubers compared to the plants with flowers/berries intact but had no impact on the average weight of tubers. There was also no effect on starch content and composition. The effect of fruit set on tuber production is generally poorly understood, as studying it requires genotypes with substantial berry production. Most commercial potato varieties produce very few berries, and therefore fruits are unlikely to represent a substantial competing sink in those lines. A previous study generated substantial variation in fruit set by crossing tetraploid lines with Phureja, but flower removal in this background had no effect on tuber yield (Kidane-Mariam & Peloquin, 1974).

Other high fruit-yielding varieties, also derived from crosses with Phureja, were subjected to a similar flower removal experiment, and this increased tuber yield in some but not all varieties, and not in all environments (Jansky & Thompson, 1990) Where total tuber yield increased, it was not determined whether the increase occurs via more tubers, larger tubers, or both. Another study found that preventing fruit formation in a heavy fruit-bearing cultivar increased tuber yield by increasing tuber size, tuber number, and dry matter content (Tekalign & Hammes, 2005). This contrasts B100, where the primary effect of fruit removal was on tuber number. However, these data were from a field experiment in Ethiopia, which represents a very different environment to our study. Thus, these contrasting findings could be due to the different environments, but also the varieties used. Since the frequency of fruit formation was not quantified in the current or previous work, it is difficult to compare the extent to which berries represent a competing sink between the different varieties tested. However, B100 is an ideal variety for study of sink-source relationships, particularly for investigating the relationship between resource allocation and tuber initiation. For example, it is known that sucrose can stimulate stolon initiation, but the underlying molecular mechanisms are unknown (Wang et al., 2025). The induction of tubers via fruit removal in B100 may represent an in vivo system where the effect of sucrose can be explored.

It is also important to note that the number of tubers produced by B26 cannot be explained by differences in resource allocation alone, as removal of flowers in B100 did not increase tuber numbers to the level of B26. This is at least partly due to flower removal promoting more vegetative growth (Figure 3C), which is consistent with previous work (Tekalign & Hammes, 2005), and prevents all assimilates from being diverted into tuber formation. However, there could be other variation in the molecular mechanism of tuber initiation in B26, leading to the high number of tubers.

### Unique genetic variation in B26 leads to novel starch

Profiling starch in these diploid lines revealed novel phenotypes that could be used to improve tuber quality. Dry matter content of tubers only varied slightly among the lines, and there were no differences in total starch content (Figure 4B). However, there were two key features of the starch structure in B26 that was different from the other lines - high resistant starch and elongated granule morphology. High resistant starch in potato can arise due to high amylose (Harris & Warren, 2024), yet our results show no significant differences in amylose content across diploid lines compared to Clearwater (Figure 4D). We did notice a trend of higher amylose content in the diploid lines, which can be attributed by the variation of measurements; however, all these values fall within the expected 20% to 33% range of amylose content in potatoes as reported in literature (Alvani et al., 2011; Kaur et al., 2007; Liu et al., 2007; Pycia et al., 2012). Therefore, we suggest that the high resistant starch content in B26 is due to other factors. Granule size also has an influence on digestibility, where larger granules digest more slowly in vitro (Dhital et al., 2010). Although B26 granules were significantly larger than those of B100, size alone cannot explain the elevated resistant starch since YD3 granules were equally as large as B26 granules, but YD3 had normal resistant starch levels (Figures 4 and 5). The elongated shape alone cannot explain the high resistant starch either, since YD2 granules had the same shape profile as B26 granules without having elevated resistant starch (Figures 4 and 6). The fact that the two hybrids inherited either the granule shape or size characteristic of B26 suggests that different genetic loci control these two traits. It is possible that other properties of the B26 starch granules, including smoothness and structural changes that form during digestion, affect digestion rates (Tian et al., 2024).

The genetic variation underpinning the unique granule shape of B26 is not known. The segregation of the elongated granule morphology phenotype among the hybrids YD2 and YD3 suggests that a single dominant heterozygous allele in B26 may be responsible in shaping the elongated granules. This could be further studied by generating F_2_ progeny from YD2, which will likely segregate for the phenotype.

Analysing more hybrids from the B26 ✕ B100 cross with quantitative granule shape analysis, such as the QICPIC (Figure 6), could allow mapping the genetic variation responsible for the phenotype. A possible candidate gene is PROTEIN TARGETING TO STARCH 2b (PTST2b), which was recently implicated in the control of granule shape in potato (Hochmuth et al., 2025). PTST2b specifically affects granule shape but not granule size, and is required for anisotropic growth of granules since reducing its expression leads to near-spherical starch granules.

In conclusion, the inbred diploid potato lines from the Solynta breeding program presented unique physiological and starch phenotypes and are excellent models for studying tuber yield and quality traits. Given that the variation that was observed in just two lines, analysing more lines is likely to reveal more valuable traits for genetic studies and breeding.

## MATERIALS AND METHODS

### Plant growth, flower disruption and tuber harvesting

Tissue culture cuttings of diploid research lines B26 and B100 from Solynta, diploid hybrids YD2 and YD3 from Yordan Dolaptchiev, and commercial tetraploid line Clearwater Russet were grown in a glasshouse from November 2023 to April 2024. Plant growth was supported with 600W LED lights with 16 h light at 20°C and 8 h dark at 16°C. Plants were grown for 17 weeks, then shoot biomass was measured using a spring scale (removing berries where present before measuring), and all tubers within the pot were harvested.

For the experiment with the removal of flowers, B100 plants in pots were distributed across the glasshouse bench, and plants for flower removal were allocated using a random block design. Flower disruption was performed by mechanically removing all parts above the calyx using tweezers: including corolla, stamens, and ovaries. The process was performed 3 times a week for each plant until harvest.

### Starch staining in berries

Berries from B100 plants were harvested from 12-week-old plants, cut longitudinally, then incubated for 5 minutes in a 1:5 solution of Lugol’s iodine solution to deionised water.

### Tuber yield, dry matter content and sampling

Tuber number per plant and total tuber weight per plant was determined for each plant after harvesting. The 3 largest tubers were sampled for each biological replicate, where the tubers were diced, mixed and lyophilised for 3 days. The weight of the samples was measured before and after lyophilisation to determine the dry matter content, expressed as the percentage of dry weight compared to the fresh weight. Freeze-dried tuber samples were ground to flour using a Krups KM75 coffee grinder and stored at room temperature for starch analyses.

### Starch purification of potato tubers

Starch granules were purified by incubating ∼300 mg of the lyophilised potato flour in 5 mL 0.5 M NaCl on ice for 30 min. Suspensions were filtered through a 100 μm nylon mesh with an additional 5 mL of water, before centrifugation for 15 min at 2500*g* at 10°C. The starch pellet was washed three times in 90% (v/v) Percoll solution (50 mM Tris-HCl, pH 8), with centrifugation for 15 min at 2500*g* at 10°C between each wash; then washed three times in 2% (w/v) SDS with centrifugation for 5 min at 3200g and room temperature between each wash; then washed three times in cold acetone with centrifugation for 5 min at 3200g and 4°C between each wash. The pellet after the final wash was air dried.

### Characterisation of starch granule morphology

For analysis of starch granules using Scanning Electron Microscopy (SEM), purified granules were dried onto a glass cover slip and mounted onto an SEM stub.

Samples were sputter coated with gold (8-9 nm) using a ACE600 sputter coater (Leica, Newcastle upon Tyne, UK) and then visualised using a Nova NanoSEM 450 (FEI, Hillsboro).

For analysis of volumetric granule size distributions, purified starch granules were analysed on the Multisizer 4e Coulter counter (Beckman Coulter), fitted with a 200 µm aperture. For each sample, approximately 5 mg of starch was suspended in 500 µL of Isoton II electrolyte solution and incubated at room temperature with mixing for 30 mins. The sample was further diluted in 100 mL electrolyte and relative volume vs. diameter data was collected for a minimum of 50,000 particles per sample.

For analysis of particle shape (aspect ratio) distributions, we used a dynamic image analysis sensor QICPIC with wet dispersion unit LIXELL (Sympatec) and an M5 optical module with a particle size optimum range of 16 to 1252 µm. For each sample, approximately 50 mg of purified starch was diluted in 150 mL of deionized water. Imaging was performed at 80 FPS for 30 seconds with an optical concentration of 0.2%. The Aspect ratio ψ_A_ (0 < ψ_A_≤ 1) is defined by the ratio of the minimum to the maximum feret diameter ψA = x_Feret min_ / x_Feret max_.

### Total, resistant starch and amylose content

Resistant and non-resistant starch fractions were assayed using the Resistant Starch Assay Kit (K-RSTAR; Megazyme). We followed the manufacturer’s protocol with minor changes by scaling down sample size to 10 mg of potato flour and adjusting reagent volumes accordingly. Total starch was calculated as the sum of both starch fractions. Amylose content was measured from purified potato starch as described by Knutson and Grove (1994) using potato type III amylose from Sigma (A0512) as standard.

### Amylopectin chain length distribution

Debranched starch samples were prepared from the starch based on the method described in Streb et al. (2008), but with modifications: 0.2 mg of purified starch was suspended in 450 µL of water and boiled for 15 min to gelatinise the starch. The starch was debranched with 0.02 U isoamylase from Pseudomonas (Megazyme) and 1 U pullulanase M1 from *Klebsiella planticola* (Megazyme) in a 500 µL reaction in 10 mM sodium acetate, pH 4.8, 37°C, 2.5 hours. The reaction was boiled for 10 min to stop the reaction. The debranched starch was purified over sequential columns of AmberChrome, before analysis using High-Performance Anion Exchange Chromatography with Pulsed Amperometric Detection (HPAEC-PAD) on a Dionex ICS-5000-PAD fitted with a PA-200 column (Thermo).

### Data and Statistical analyses

Data analysis for tuber and plant traits consists of tuber number, total tuber weight per plant, shoot biomass, fresh and dry matter percentage, and average tuber weight where total tuber weight of each biological replicate was divided by its corresponding tuber number. Data analysis for starch properties includes total starch content, with resistant starch and non-resistant starch fractions, and mean granule size. Line plots for Coulter counter and QICPIC show smoothed data using a rolling average of 5 data points.

Data representation is divided in 2 groups: Group 1 showing data from all 5 lines, where B100 data is of untreated WT replicates, and Group 2 comparing untreated versus treated replicates for B100 line. All statistical analyses for Group 1 were done using One-Way ANOVA, while Group 2 analyses were done using Student’s t-test. Figures and data analysis were performed in R version 4.4.1.

## ACKNOWLEDGEMENTS

We thank Solynta for providing access to B26 and B100 lines, John Innes Centre (JIC) Horticultural Services for providing growth facilities, Phil Robinson (JIC) for providing photography, Fred Warren (Quadram Institute) for providing access to the QICPIC instrument, Rose McNelly (JIC) for assisting with QICPIC analysis, and Brendan Fahy (JIC) for advising with the amylose content assay.

## DECLARATIONS

### Availability of data and materials

All data generated during this study are included in this article and its supplementary information files.

## Authors’ contributions

**Thomas Navarro:** designed and conducted experiments, analysed data, generated figures, wrote, reviewed and edited original draft. **Yordan Dolaptchiev:** provided diploid material, consulting, reviewed and edited final draft. **Oluwaseyi Bello:** conducted experiments, analysed data, reviewed and edited final draft. **Ciara**

**O’Brien:** conducted experiments, analysed data, consulting, reviewed and edited final draft. **Andrés Ortiz:** conducted experiments, analysed data, reviewed and edited final draft. **Jonathan Jones:** designed experiments, consulting, supervision, reviewed and edited final draft. **David Seung:** designed experiments, consulting, supervision, wrote, reviewed, and edited original draft.

## SUPPLEMENTARY MATERIAL

**Figure S1.**
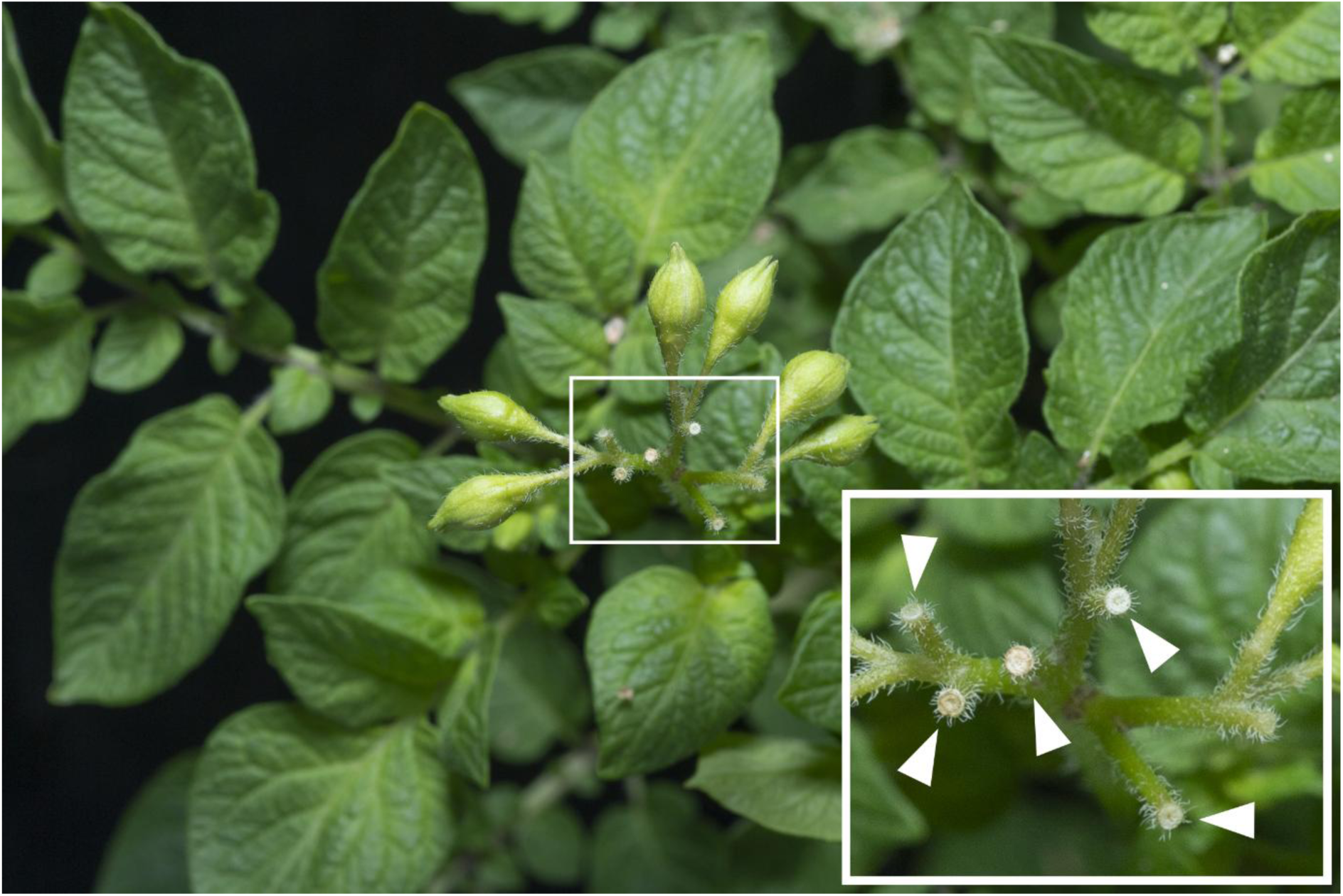
**Flower abortion phenotype of B26**. Photograph of flower buds from diploid line B26. Bud abscissions are marked with arrows.

**Figure S2:**
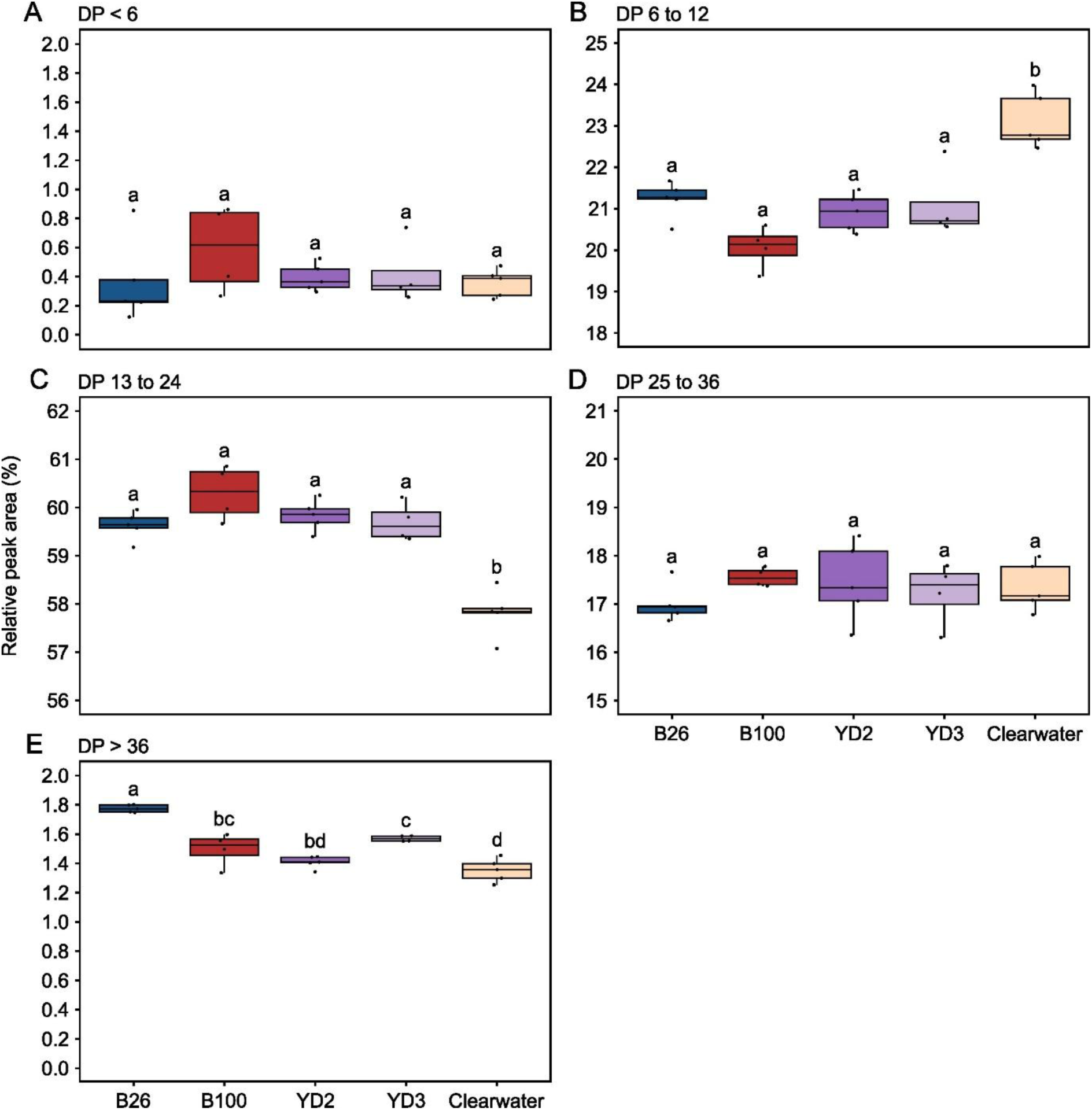
Diploids have an altered amylopectin chain length distribution compared to Clearwater Russet. Analysis of the amylopectin chain length distribution from Figure 4E by averaging degree of polymerisation (DP) values in five representative groups: **(A)** Chains smaller than 6 glucose units, **(B)** chains between 6 and 12 glucose units, **(C)** chains between 13 and 24 glucose units, **(D)** chains between 25 and 36 units, and **(E)** chains longer than 36 glucose units. Values represent the total relative peak area as a percentage for each group. Dots represent individual data points (n = 4-5 per genotype, with each point representing tubers harvested from an individual plant). For all boxplots in this figure, the band inside the box represents the median, while the top and the bottom of the box represent the lower and upper quartiles, respectively. The whiskers represent values within 1.5× of the interquartile range. Values with different letters are significantly different from each other under a one-way ANOVA with Tukey’s posthoc test (p <0.05).

